# Detect+Track: Robust and flexible software tools for improved tracking and behavioural analysis of fish

**DOI:** 10.1101/2023.04.20.537633

**Authors:** Abhishek Dutta, Natalia Pérez-Campanero, Graham K. Taylor, Andrew Zisserman, Cait Newport

## Abstract

Developments in automated animal behavioural analysis software are increasing the efficiency of data collection and improving the standardization of behavioural measurements. There are now several open-source tools for tracking laboratory animals, but often these are only accurate under limited conditions (*e*.*g*. uniform lighting and background, uncluttered scenes, unobstructed focal animal). Tracking fish presents a particular challenge for these tools because movement at the water’s surface introduces significant noise. Partial occlusion of the focal animal can also be troublesome, particularly when tracking the whole organism. But identifying the position of an animal is only part of the task – analysing the movement of the animal relative to their environment and experimental context is often what provides information about their behaviour. Therefore, the automated detection of physical objects and boundaries would also be beneficial, but this feature is not commonly incorporated into existing tracking software. Here we describe a video processing method that uses a range of computer vision algorithms (*e*.*g*. object detector and tracker, optical flow, parallel plane homology) and computational geometry techniques (*e*.*g*. Voronoi tessellation) to analyse the movement behaviour of fish in response to experimental stimuli. A behavioural experiment, which involved tracking a fish’s trajectory through a field of obstacles, motivated our development of a set of tools that: (1) measure an animal’s trajectory, (2) record obstacle position, and (3) detect when the fish passed through ‘virtual gates’ between adjacent obstacles and/or the aquarium wall. We have introduced a novel Detect+Track approach that significantly enhances the accuracy and robustness of animal tracking, overcoming some of the limitations of existing tools and providing a more reliable solution for complex experimental conditions. Our workflow is divided into several discrete steps, and provides a set of modular software building blocks that can be adapted to analyse other experimental designs. A detailed tutorial is provided, together with all the data and code required to reproduce our results.

---

For many, an aquarium may be a home for fish or a work of art, but for a researcher, an aquarium can serve as a self-contained laboratory. Increasingly, behavioural experiments are using fish as models to answer diverse questions over a range of fields including neuroscience (***Steward et al., 2014***; ***Orger and de Oilavieja, 2017***; ***Roberts et al., 2022***), medicine (***Shenoy et al., 2022***), sensory ecology (***Marshall et al., 2018***; ***Newport and Schuster, 2020***), cognition (***Pouca and Brown, 2017***), biomechanics (***Liao, 2007***), and bio-inspired robotics (***Hu et al., 2006***). In many cases, the behavioural metrics of interest relate to spatiotemporal patterns of movement measured under different experimental conditions. These include distance travelled, speed of movement, position relative to a goal or stimulus, and quantification of exploratory behaviour. It is now common to extract these data automatically from experimental video recordings, which has the benefit not only of being faster, but – depending on the accuracy of the tracking system – also of increasing the accuracy and repeatability of the measurements. For instance, subjective identification of a single point representing the animal’s position is observer dependent and variable, but computer vision can turn this into a standardized mathematical process of identifying the animal’s centroid.

Some of the most accessible software solutions for laboratory animal tracking use background subtraction (*e*.*g*. ToxTrac ***Rodriguez et al. (2018***), TRex ***Walter and Couzin (2021a***)) or blob detection (*e*.*g*. idTracker ***Pérez-Escudero et al. (2014***)). Object detection using these methods has a relatively fast processing time, and does not require any additional information other than the video of interest. For example, in background subtraction, some or all of the video frames are averaged to create a reference image: every feature that is present and in the same position throughout the video will appear in this background image. When a frame that includes a moving animal is compared to this background, pixel areas which differ substantially from the background are identified as foreground. These foreground pixels ostensibly indicate the position of the animal if it is the only moving foreground object, but this will rarely be the case in natural settings. Moreover, this method will detect anything that moves, and when filming animals through an air-water interface, many regions can move. Ripples and reflections in particular will often cause a fish to be lost amongst all the other moving pixels. However, with careful camera positioning ensuring limited reflections, and tuning of image characteristics (*e*.*g*. pixel region or blob sizes, image contrast) this method can still be effective for experimental applications, even when image clarity is low (*e*.*g*. ***Newport et al. (2021***)).

Where background subtraction fails, more sophisticated computer vision approaches can be used. For standard fish models, such as zebrafish (*Danio rerio*), there is already bespoke software capable of tracking fish under common experimental conditions (***Franco-Restrepo et al., 2019***). However, tracking accuracy can be low, or may fail altogether, when applied to different experimental scenarios or species. Often the reason for the failure is not because the task is impossible, or even particularly challenging by current computer vision standards, but rather because the underlying tracking method is not appropriate to the task. Advanced machine learning approaches provide a general solution, but for many biologists, the steep initial learning curve associated with applying these techniques still represents a barrier to entry. A deep learning approach to fish detection and tracking (***Hunt et al., 2021***; ***Lauer et al., 2022***) can produce more robust results under a wider range of conditions including changes in lighting throughout the video, partial occlusion of the focal animal, the presence of multiple individuals, and interference from significant surface ripples. However, deep learning approaches are best-suited to feature tracking, and whilst this is often desirable (*e*.*g*. in pose estimation), there may be other instances where it is necessary to track the centroid of a deforming object (*e*.*g*. in biomechanical applications tracking the centre of mass).

Once the position of an animal has been identified, computer vision can also be used to characterize or quantify other relevant behaviours. For example, changes in the speed or direction of an animal throughout the experimental period, along with information about its movement within the experimental area and in relation to experimental stimuli, can serve as important behavioural indicators. Determining an animals proximity to edges or experimental walls, are not only useful to know in terms of when they have reached a movement limit, but can also be useful to explore the time spent near experimental edges or behaviours such as thigmotaxis (orientation using physical contact). Identifying time spent in different tank areas, or an animals proximity to important stimuli are commonly used in tests of stimuli preference and avoidance.

Here, we develop a workflow that combines deep learning with classical computer vision techniques to improve fish tracking accuracy in a non-standard experimental setting that includes surface ripples as well as object stimuli that partially occlude the focal animal. In addition to providing accurate centroid tracking of the focal animal, our workflow also analyses fish movement behaviour (*i*.*e*. speed, movement direction), as well as key features of the experimental setup (*i*.*e*. obstacle presence and position) to which it relates the animal’s motion (*i*.*e*. gap negotiation between obstacles).

## Results

In our experiment, we filmed *n* = 5 Picasso triggerfish (*Rhinecanthus aculeatus*) from above the water’s surface, as they moved individually through a field of cylindrical obstacles within an aquarium to reach a food reward (Figure 1). The goals of this behavioural experiment required us: (i) to identify the path taken by the fish; (ii) to determine which of the gaps between obstacles the fish selected; and (iii) to record the size of each gap. A trial ended once the fish had reached the food reward, so identification of this location was also required.

**Figure 1.**
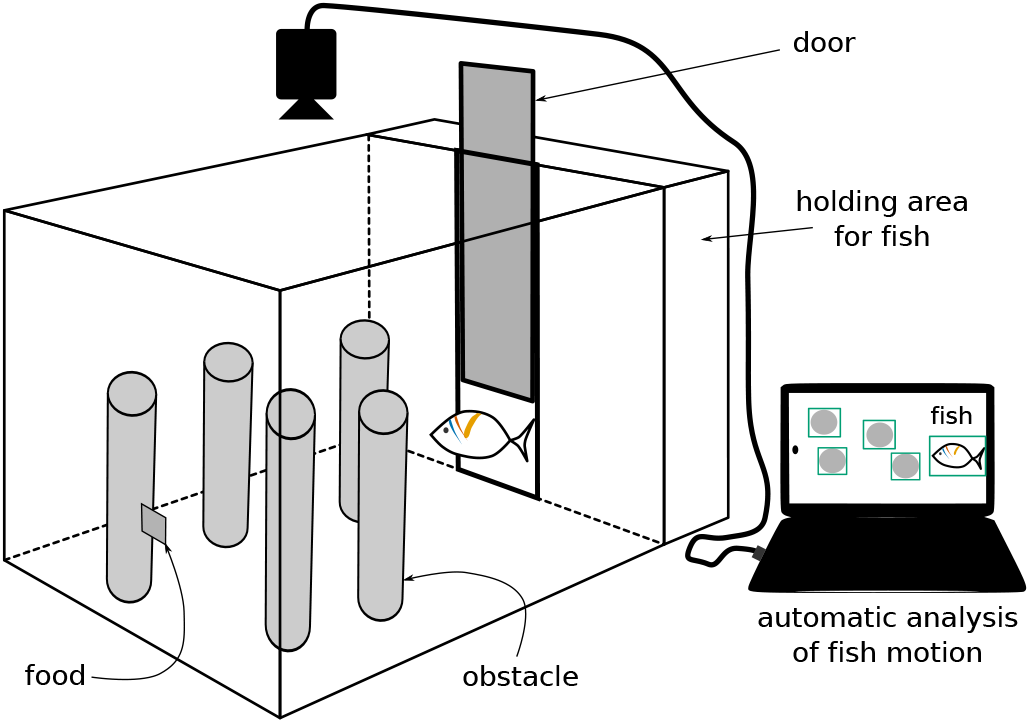
A experimental aquarium containing fish, obstacles and food is monitored by an overhead camera. The resulting video is analysed by the automated processing workflow described in this paper.

We used the experimental setup shown in Figure 1 to capture 454 videos showing the journey of a fish. A set of 20 randomly selected videos – called the training set – were used to train our deep neural network based fish and obstacle detectors. Another randomly selected sample of 15 videos – called the evaluation set – were used to evaluate the performance of our workflow. As a result of this sampling, the fish detector was trained using videos from three fish, while the evaluation set included all five fish. The remaining 419 videos were automatically processed using our workflow as illustrated in Figure 2.

**Figure 2.**
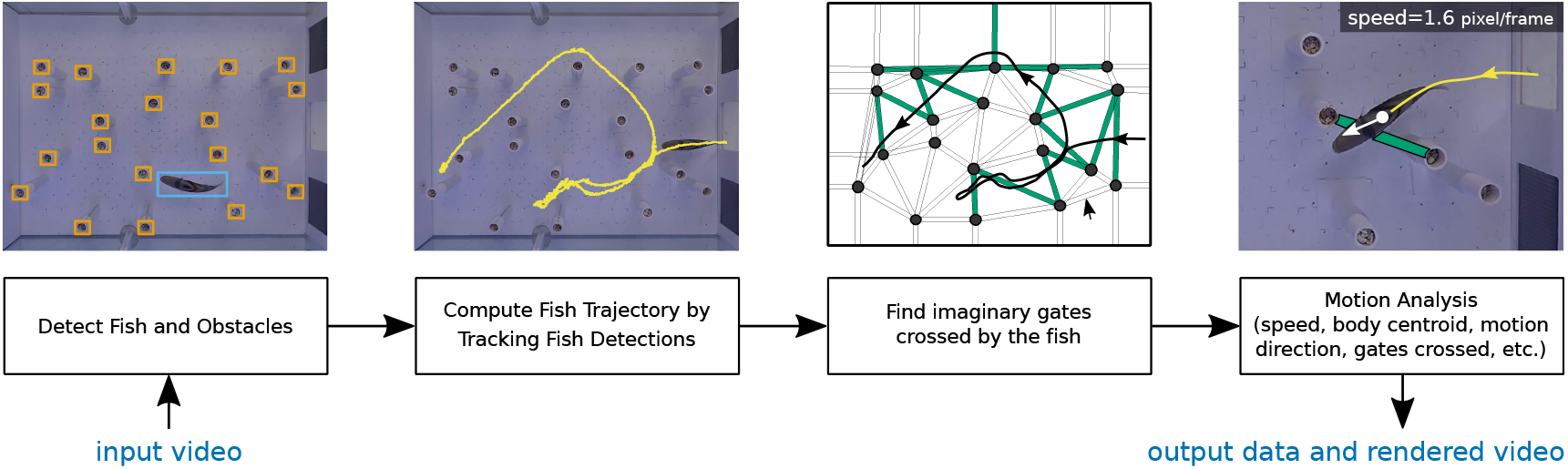
Illustration of the automatic video processing method described in this paper. Frames are extracted from the input video and fed into an object detector that can detect the location of the fish and the obstacles. Next, an object tracker is applied to high-confidence fish detections in order to build the fish motion trajectory. Virtual gates formed by obstacles are located next based on automatic detection of cylindrical obstacles and calibration feature points. The virtual gates and fish trajectory help locate all the gaps between obstacles that the fish passes through during its journey. Finally, fish motion is analysed using a dense optical flow method to estimate speed, motion direction and body centroid.

Each video in our dataset represents an individual behavioural trial. The location of the fish in each video was computed using a “Detect+Track” method which operates in two stages. In stage one, an object detector (*i*.*e*. fish detector based on EfficientDet ***Tan et al. (2020***)) is applied to automatically detect fish with high confidence only in some frames of a video. In stage two, an object tracker capable of robustly tracking any object over a short interval is applied to fill in the gaps. This two stage Detect+Track method results in precise localization of the fish centroid compared to ground truth (i.e. more accurate) and consistently detects the fish (i.e. more robust). The fish detector was trained using 235 manually annotated frames from the training set. The detector produced high-confidence detections (*i*.*e*. detections with > 70% confidence) in 39.8% of the frames. An object tracker, initialised with these high-confidence detections, filled in the missing detections in the intermediate frames.

### Comparison to baseline methods

We evaluated and compared our Detect+Track approach against two other commonly used software tools: idtracker.ai ***Pérez-Escudero et al. (2014***) (version 5.2.2.1, Oct 2023), TRex ***Walter and Couzin (2021b***) (version 1.1.6, Jan 2022). We used the Percentage of Correct Keypoints (PCK) metric to evaluate the accuracy of predicted body centroids compared to the manually annotated centroids in the ground truth. The PCK^1^ metric corresponds to the proportion of video frames for which the computed centroid lies within certain threshold distance (*e*.*g*. a threshold of 30 pixels) of the manually annotated fish body centroid. Because bounding boxes can change in size from frame to frame due to differences in fish size and body position, we normalised our performance metric by dividing the threshold distance (in pixels) by the square root of the fish bounding box area. We use the term *characteristic length* to refer to the square root of the area of the fish bounding box. Figure 3 shows the PCK metric at various normalized threshold values for fish body centroid obtained using our Detect+Track method and that using the idtracker.ai and TRex software tools.

**Figure 3.**
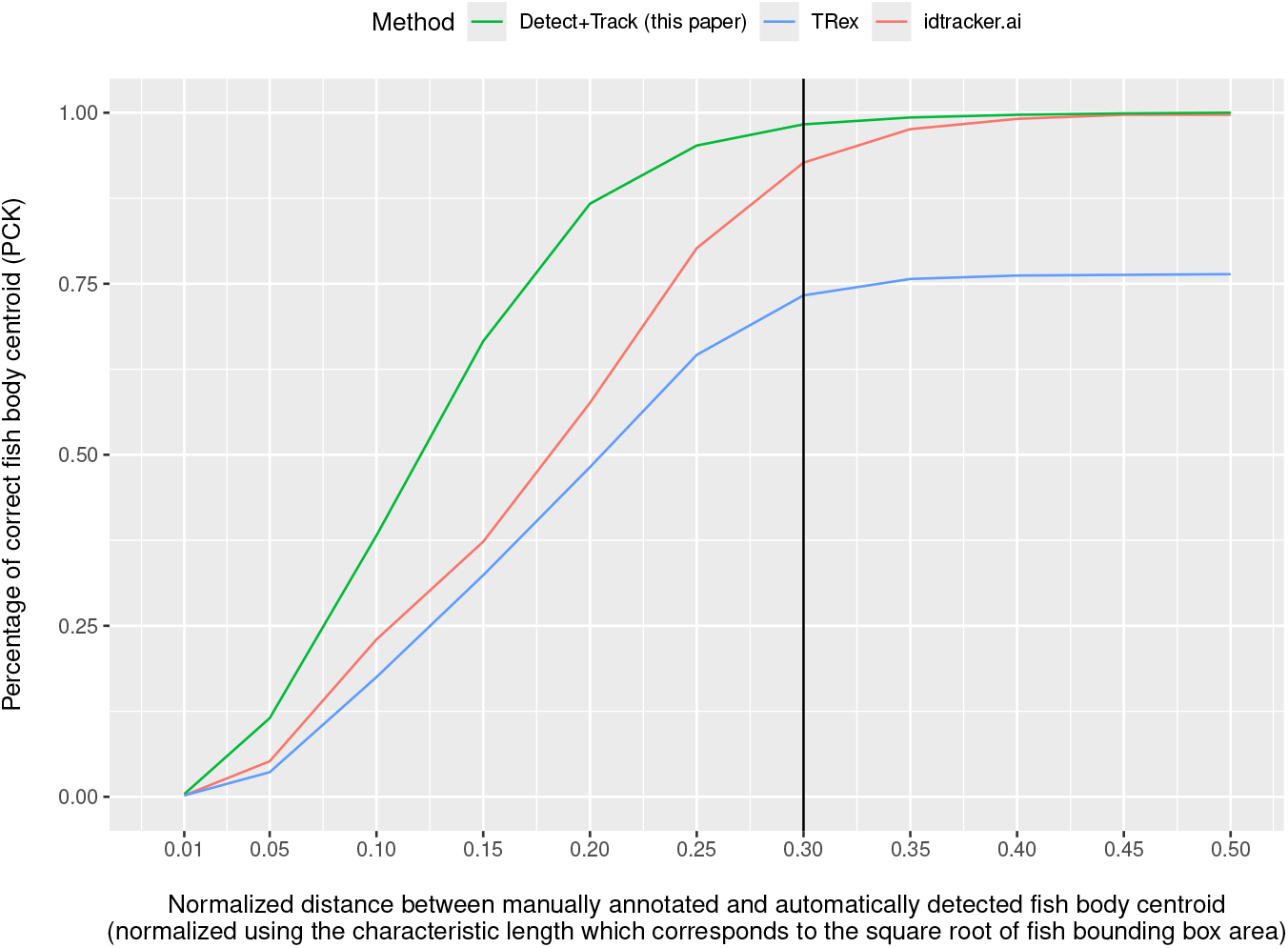
Performance comparison with existing animal tracking software tools: The accuracy of automatically computed fish body centroid is computed by measuring the proportion of video frames for which the computed centroid point lies within certain normalized distance from the manually annotated ground truth. This plot shows that our Detect+Track method (green) is more accurate and robust at estimating fish body centroid as compared to TRex (blue) ***Walter and Couzin (2021b***), and idtracker.ai (red) ***Pérez-Escudero et al. (2014***). The line at 0.3 indicates the point at which we compare the accuracy of the three methods.

The fish body centroid detected by our Detect+Track method is more accurate than that of detected by idtracker.ai and TRex. For example, 98.3% of the fish body centroids obtained by our method lie within a characteristic length of 0.3, whereas only 92.7% of the centroids obtained by idtracker.ai and 73.3% from TRex fall within this area (Figure 3). The idtracker.ai method is able to eventually reach the performance level achieved by our Detect+Track method but at the cost of higher error in estimated fish body centroid. The TRex method’s performance asymptotes at 0.75 which indicates that it is unable to improve its accuracy any further. Our Detect+Track method is also more robust at detecting fish body centroid because it is able to successfully detect fish in all frames (Table 1) while the idtracker.ai and TRex tools failed to detect fish in 98 and 105 frames, respectively of the total 3846 frames extracted from 15 videos in the evaluation subset. Detection failure in idtracker.ai and TRex typically occurred due to factors such as objects moving in the background scene (outside the tank) and ripples on the water surface.

**Table 1.**
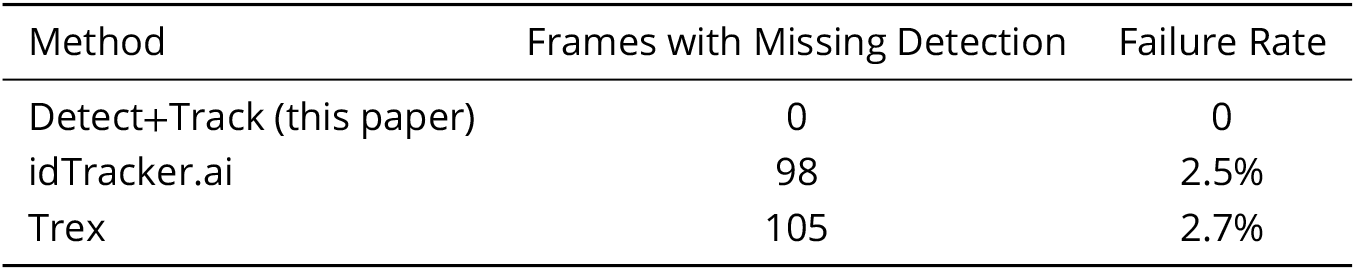
Fish detection robustness: Number of missed detection’s out of 3846 video frames for for the Detect+Track method described in this paper, TRex ***Walter and Couzin (2021b***), and idtracker.ai ***Pérez-Escudero et al. (2014***).

The goal of our behavioural experiment was to identify which gaps between adjacent obstacles were chosen by the fish during its journey to the food, in order to understand the mechanisms determining gap choice. Each video contained either 12 or 20 obstacles arranged in different spatial configurations. An obstacle detector was used to find the position of these obstacles in each video automatically. The performance of the obstacle detector was evaluated using the evaluation set in which we manually annotated the position of the obstacles. The obstacle detector was able to detect all 292 obstacles (*i*.*e*. no false negatives), with only 4 false positives. The workflow was designed so that any errors in automatic detection could be corrected by a human operator.

To identify which obstacles the fish passed between, we used Voronoi tesselation to locate each of the detected obstacles within a polygonal cell. We identified adjacent obstacles as those detected obstacles whose Voronoi cells shared a common edge, drawing a ‘virtual gate’ between them, which we took to represent the gaps between obstacles.

Finally, we used the classical optical flow formulation to estimate the fish’s apparent motion between two consecutive frames. This allowed us to compute the trajectory of the fish automatically (Figure 4), by identifying the centroid of the subset of pixels having an optical flow magnitude greater than some specified threshold. We then averaged the optical flow over these pixels in order to determine the apparent speed and direction of motion of the fish in every frame. Figure 5 shows the computed fish trajectory with coloured dots indicating the speed at each instant and an arrow to show the fish motion direction. Highlighted Voronoi cells indicate those cells that the fish visited.

**Figure 4.**
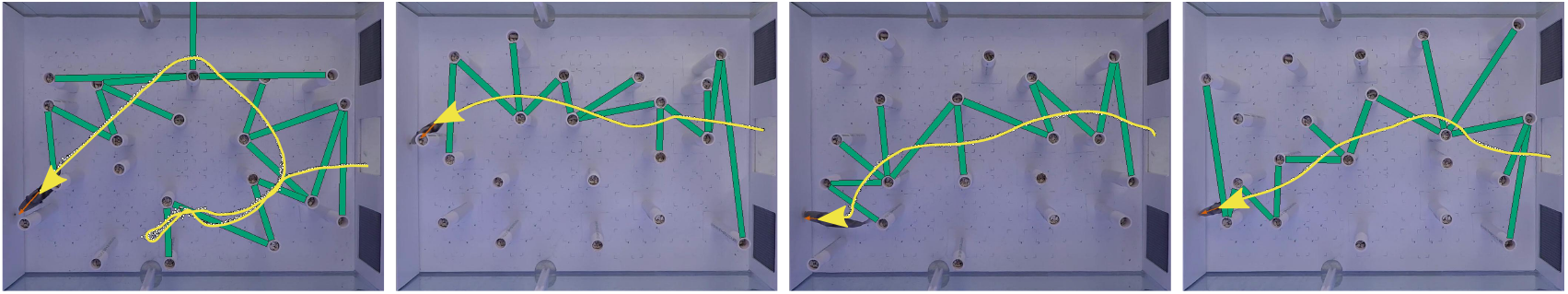
Examples of fish trajectory automatically computed by the fish video processing method described in this paper. The position of the fish through time is shown as a yellow line. The virtual gates crossed by the fish are shown in green. The four trajectories correspond to four different Picasso triggerfish subjects experiencing four different obstacle arrangements. The frames capture the instant when the fish reaches the food at the left end of the aquarium.

**Figure 5.**
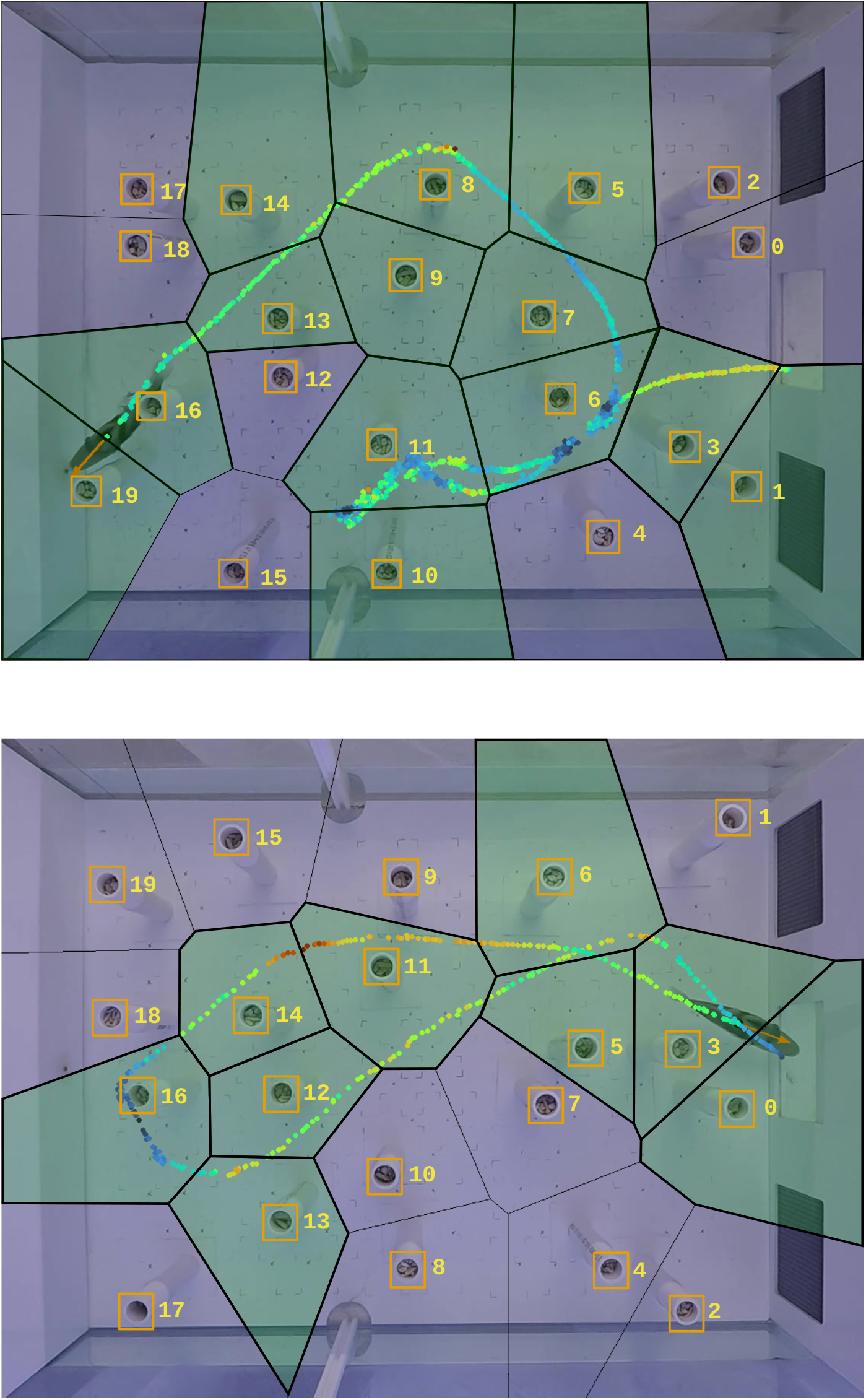
Detailed view of two automatically computed fish trajectories shown as coloured dots representing fish speed. Here, red implies high speed, while blue is low speed, with green corresponding to intermediate speed. The orange arrow on the fish shows its instantaneous motion direction. The obstacles are numbered and marked using orange rectangles. Each polygon is a Voronoi cell centered on an obstacle; green-shaded cells represent those the fish had visited.

**Figure 6.**
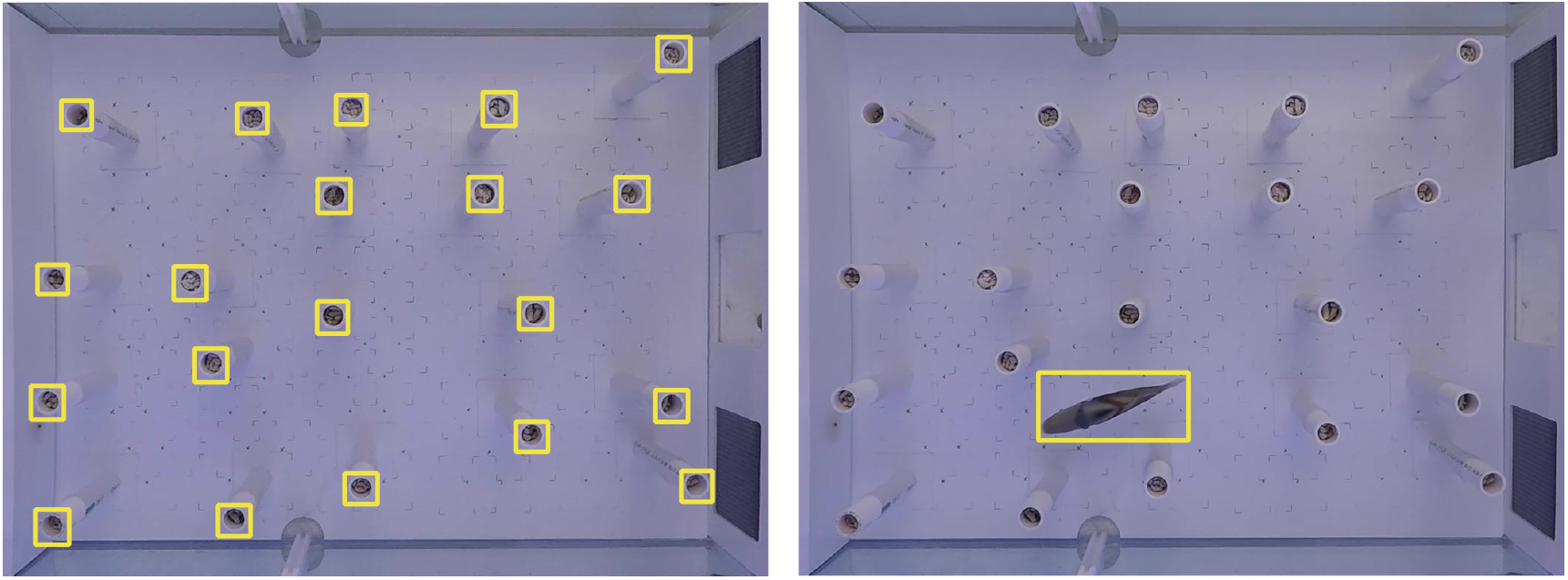
Objects of interest (*i*.*e*. fish and top part of obstacles) are manually annotated with a rectangular bounding box (yellow). These annotations are used for training the object detectors.

The software code released with this paper includes a script that allows for batch processing of videos. After the initial calibration, all 454 videos were processed automatically using this script. A 10 second video takes approximately 4 minutes to process on a 16 core 3.60GHz machine with an NVIDIA GeForce GTX 1080 Ti (12Gb) GPU. Therefore it is practicable for processing the large datasets that are often collected in behavioural assays.

## Discussion

In this paper, we have developed and described a video processing method to automatically analyse the motion a moving subject, demonstrated using the example of a Picasso triggerfish swimming in an aquarium setting. This method uses a deep-learning based object detector and object tracker combined with an optical flow estimation method to accurately identify the fish trajectory throughout an experimental trial. Our videos show an experiment in which a single fish swims through a field of obstacles in order to reach a goal. The fish changes shape as its body curves during swimming, and in some instances it is partially occluded by obstacles. In addition, the angular field of view of the camera was varied between some of the trials, which changes the apparent shape and size of the objects. These variations make the automatic detection and tracking more challenging. Our tests on the evaluation set show that 97% of the computed trajectories have more than 50% overlap (*i*.*e*. intersection-over-union) with the ground truth identified by a human expert. We also developed methods to automatically detect the location of visual stimuli (*e*.*g*. cylindrical obstacles) and identify the ‘virtual gates’ defining the gaps between object stimuli which the fish must pass through. As virtual gates are also formed between obstacles and the wall of the aquarium, a method was developed to create imaginary obstacles along a plane, and to then create gates between the real obstacle and nearest imaginary obstacle. The automatically computed fish trajectory and location of the visual stimuli gives a unique insight into the influence of deliberately placed obstacles on the behaviour of the fish.

An automatic video processing system based on computer vision will always have some cases where the automatic analysis may fail. For example, an obstacle detector may fail to detect all obstacles, or may wrongly detect their presence at a location where none exists. Our workflow supports manual correction of any failure in automatic processing by a human operator if desired. This capability to intervene manually when required allows our method to be applicable to many challenging experimental conditions, while still ensuring high accuracy. Perhaps most importantly, by combining classical computer vision techniques with deep learning, our method provides accurate information on the motion of the centroid of a deforming object, rather than some unique feature thereof. This is relevant to biomechanical applications in which it is necessary to track the motion of the centre of mass, whose anatomical position may vary as a result of body deformation or changes in pose.

The video processing method presented in this paper has some limitations and practical considerations. First, the workflow is intended to process videos containing a single animal and it would require additional tools to extend it to multiple animals. Other existing tools (***Walter and Couzin, 2021a***; ***Lauer et al., 2022***) support tracking of multiple animals in the same video. Second, the obstacles near the aquarium boundary can often cause partial occlusion of the fish which can have detrimental effect on the automatic detection and tracking. This can lead to more measurement error in obstacles near the boundary as compared to the measurements related to obstacles in the center. Finally, the object detector re-training stage requires a GPU with more than 8GB of memory. We provide pre-trained models for fish and obstacle detector which can be directly applied to other datasets by a user that lack access to such computing resources. Retraining for detecting other categories of animals is possible using the software tools released with this paper.

We have released all the assets (*e*.*g*. software, scripts, tools and videos) associated with this project under a Creative Commons license which allows unrestricted use and extension of these assets for any purpose. The goal is to enable other researchers to reuse and extend these video processing methods to suit their own requirements. These assets are also a valuable tool for biologists who want to learn and understand how modern deep learning methods and existing tools are used to collectively form a complete video processing pipeline. We expect the set of 454 videos released as a part of this paper to be useful to computer vision researchers who are exploring challenging test data sets in animal tracking. Our dataset presents a challenging scenario in which the cylindrical obstacles can occlude the fish and the fish itself can change its overall shape throughout the video. In the following section we suggest some alternative workflow applications for those wanting to assess the usefulness of our tools in relation to their own video data.

### Adapting to New Experimental Settings

Each processing stage in our workflow is implemented as a script which feeds on the output generated by previous processing stages and produces results that can be used as the input to the next stage. This modular design allows the components to be reused and extended to suit novel applications. The software code released with this paper includes a comprehensive tutorial to guide readers through all the steps involved. This tutorial can also be used as a practical introduction to application of computer vision tools in behavioural experiments involving detection and tracking of animals and stimuli, and is designed to be readable and intelligible for a wide range of audiences. The behavioural experiments described in this paper involve Picasso triggerfish and several white cylindrical objects, and the code repository accompanying this paper contains pre-trained object detectors to automatically detect both. We also provide software tools and instructions for training object detectors for detecting new focal species or experimental stimuli following the method described in Section Automatic Detection of Fish and Obstacles. The EfficientDet object detector used in our workflow can be trained to detect any object (*e*.*g*. mouse, chimpanzee, bees, *etc*.) but the detector’s performance may drop if the focal object is very small or well camouflaged. The object detector has proved to be highly accurate in our experiments where a relatively large fish (total length: Fish 38I = 165mm, fish 39J = 155mm, fish 40M = 130mm, fish 41N = 126mm, Fish 30 = 128mm) was filmed within a relatively small field of view (600 × 740 mm). The tracker used in our workflow to fill in any object detection gaps, is applicable to any object and does not require training.

Automatic identification of stimuli is a requirement in many behavioural experiments. For example, the location of experimental stimuli, food rewards, refuge areas, *etc*., are often important. Where objects are visually distinct, they can be automatically detected. However, our workflow also includes manual annotation tools that allow human operators to identify the location of stimuli which can not be easily detected automatically (*e*.*g*. very small objects such as a food target, objects that can vary dramatically in appearance, or those that are partially occluded, *etc*.).

We apply the homology of parallel planes to compute the location of virtual gates formed at the boundary, as described in Section Locating virtual gates Formed by the Obstacles. This formulation can be used in any other application where one needs to identify points in one plane that correspond to points defined either manually or automatically in another plane. For example, the formulation can be used to find the height of an object based on the known height of other objects within the scene.

The optical flow formulation applied to consecutive video frames allows us to analyse the fish motion and to compute its body centroid, speed and motion direction. Such optical flow based measurements can be applied to other experimental settings as well without changes; the only requirement being that one has identified the location of the focal animal in consecutive video frames.

The final result of our workflow is a JSON text file containing the following details:

- spatial coordinates of the obstacles and of the virtual gates formed by the obstacles, as well as the size (or length in pixels) of these gates,
- Voronoi tesselation of the fish tank area, where each tile contains an obstacle,
- spatial coordinates of the bounding box around fish body as well as coordinates of fish body centroid, motion vector and speed (in pixels/second) in each frame,
- a list of all virtual gates crossed by the fish along with the time of crossing.

This text file can be imported using other software (*e*.*g*. R, Matlab, Python, *etc*.) to pursue further advanced analysis of fish motion. For example, in addition to identifying which gate the fish went through, we could also examine the time spent in different areas of the tank defined by the Voronoi tessellations. Alternatively, the crossing of virtual gates could be applied to experimental scenarios where an animal must pass into a response area (*e*.*g*. T-maze arms).

## Materials and methods

### Behavioural Experiments

We used Picasso triggerfish *Rhinecanthus aculeatus* as the focal animals in this experiment. This coral reef species has been used previously in studies of visual ecology (***Cheney et al., 2022***; ***Newport et al., 2021***; ***Matchette et al., 2020***), and is well suited to laboratory experiments. All experimental and husbandry protocols were approved by the Animal Welfare and Ethical Review Board of the University of Oxford, Department of Biology, and methods were carried out in accordance with relevant regulations and the code of practice for the care and use of animals for scientific purposes. A total of *n* = 5 wild-caught fish were used in the experiments, purchased from local suppliers. They were held in individual aquariums (36×104cm) in a flow-through marine aquarium system and were fed a mixed diet of pellets (Ocean Nutrition, Formula One Marine Pellet) and fish, squid, prawn and cockles.

During experiments, fish were moved from their home aquarium to the experimental tank (60×80cm) by hand-net. To attach a food reward to the aquarium wall, a single pellet was set within agar agar jelly in a small plastic petri dish (see ***Newport et al. (2021***) for preparation technique). This was attached to the wall by hook and loop fasteners. Depending on the trial, either 12 or 20 obstacles were placed within the experimental area. Fish participated in 5-10 consecutive trials per day and were then returned to their home tank. Individual fish completed between 85-94 trials in total.

Trials were filmed from above by a GoPro Hero 5 camera mounted over the aquarium (resolution: 1080p, frames per second: 60, field of view setting: ‘linear’ or ‘wide’). As we recorded one continuous video each day, the videos were cut into trial sections manually prior to automatic processing, using Imaging Studio 4. Files were initially saved as ‘.avi’ (Raw, uncompressed) but were later converted to ‘.mp4’ (compressed using ffmpeg tool with libx264 video codec and crf of 23).

### Automatic Detection of Fish and Obstacles

Locating objects of interest in an image is a general requirement for many practical applications. Recent advancements in deep learning have enabled the development of a family of neural networks, called *object detectors*, that can accurately and automatically detect the location of common objects within an image. Existing object detectors can detect and recognise common object categories, but can also be easily re-trained to detect novel object categories. This capability to learn new objects has enabled the widespread application of detectors to a range of tasks including detection of illustrations in early printed books (***Dutta et al., 2021***), chimpanzees in the wild (***Bain et al., 2021***), and the position of analog clocks in photographs (***Yang et al., 2022***). Here we have re-trained object detectors to automatically detect Picasso triggerfish and cylindrical obstacles.

Samples of an object’s appearance as seen in photographs or frames extracted from a video are required to re-train an existing object detector. A set of 20 video recordings of the fish (*i*.*e*. the training set) were used for training the object detectors. We manually annotated the fish using rectangular bounding boxes in 235 frames sampled at 1 second intervals. To increase our training set, an object tracker was then used to fill in the fish annotations for the intermediate frames. The results were then manually verified, and corrected if needed, to create a training dataset consisting of 5665 annotated fish. Because the obstacles remain static throughout a video, they are only annotated in the first frame of each video (total 360 obstacle annotations in a frame of 20 videos each containing either 12 or 20 obstacles). Annotations were made using the List Annotator (LISA) tool developed by ***Dutta and Zisserman (2019***).

The size of the training data set for the obstacle object category is artificially increased – a standard practice in deep neural network training called data augmentation – by generating new images from existing video frames. For example, rotating the frames by 180◦generates a new view of the fish tank with a new spatial arrangement of the obstacles. Such generated training samples helps the object detector retain its performance when applied to new videos with a new spatial arrangement of the obstacles. Another way of generating a new image is to swap pixels values between the left and right (or top and bottom) halves of the image. This operation is often called horizontal (or vertical) pixel flip operation. All of these methods are also applied to the fish object category in order to synthetically increase the size of the training data set.

We chose to use and re-train the EfficientDet objector detector developed by ***Tan et al. (2020***) because it can easily be trained to achieve high accuracy using a small number of training examples. This object detector has already been trained to detect 80 common object categories including bus, apple, television, *etc*. Most of the training parameters are well documented and set to their sensible default values which allows one to obtain a fairly accurate object detector without requiring research expertise in the field of deep learning.

### Detect+Track: Tracking High-Confidence Fish Detections

The automatic fish detector, described above, can detect fish with high confidence^2^ only in certain video frames. Therefore, the location of the fish remains unknown in many intermediate video frames. These missing detections result in a sequence of consecutive frames without any fish detection. By definition, any such missing segment is preceded and succeeded by a high-confidence fish detection. We use an object tracker to fill in these missing detections. An object tracker can follow any object in a video after being initialised with a template image of the object in one of the initial frames. The object tracker achieves its tracking objective by assuming that the object of interest can only move a few pixels in one of the possible directions in consecutive frames of a video. It uses a template matching strategy (*e*.*g*. cross correlation) to find the new position of the object in adjacent video frames. We use the deep neural network based object tracker introduced by ***Li et al. (2018***). We did not have to train this object tracker as a pre-trained^3^ model of the object tracker is already available. We use the term “Detect+Track” to refer to this approach.

This template based object tracker can be initialised using a template in one of the video frames and can be applied either forward (*i*.*e*. next video frame) or backward (*i*.*e*. previous video frame). For forward tracking, the template based object tracker is initialised with the high confidence detection available immediately before the missing segment and allowed to track forward in time until the mid point of the missing segment. In a similar way, for backward tracking, the tracker is initialised with the high confidence detection available immediately after the missing segment and is allowed to run backward in time until the tracker reaches the frame that lies at the center of the missing segment. This process, illustrated in Figure 7, generates an accurate trajectory of the fish because the tracking process is initialised with high confidence detection and needs to be applied only for the few frames within the missing segment.

**Figure 7.**
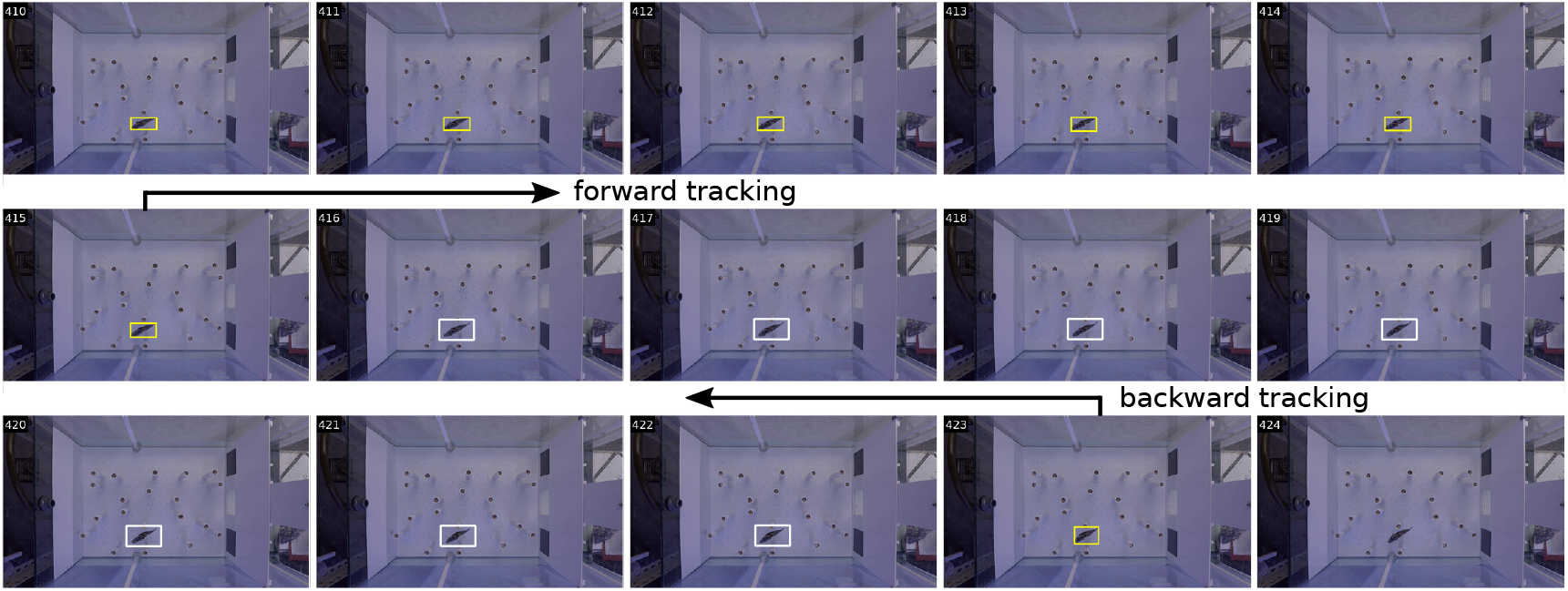
An object tracker is used to fill detections for a missing segment (*i*.*e*. a sequence of frames without any fish detection). The object tracker is initialised with high confidence detection (shown as yellow bounding boxes) available before the missing segment and is allowed to run forward in time (*i*.*e*. forward tracking) until it reaches the mid point of the missing segment. Similarly, the object tracker is initialised with high confidence detection available after the missing segment and is allowed to run backward in time (*i*.*e*. backward tracking) until it reaches the mid point of the missing segment. Such forward and backward tracking is used to locate fish (shown as white bounding boxes) in all the missing segments.

### Locating virtual gates Formed by the Obstacles

A fish crosses multiple virtual gates, defined in Box 1, before reaching the food target. In this section, we describe our method for computing the location of these virtual gates, which we categorise as either obstacle gates formed between adjacent obstacles, or as boundary gates formed between an obstacle and the aquarium wall.

The Voronoi cell tessellation method described in ***O’Rourke (1994***) is applied to find the location of all obstacle gates. Each Voronoi cell contains an obstacle and all points within a cell are closer to the contained obstacle than to any other obstacle. The boundaries of the Voronoi cells are approximated as polygons (Figure 8a). Obstacles are said to be adjacent if their Voronoi cells share an edge in common, such that the line connecting a pair of adjacent obstacles defines the virtual gate between them (Figure 8a; orange lines).

**Figure 8.**
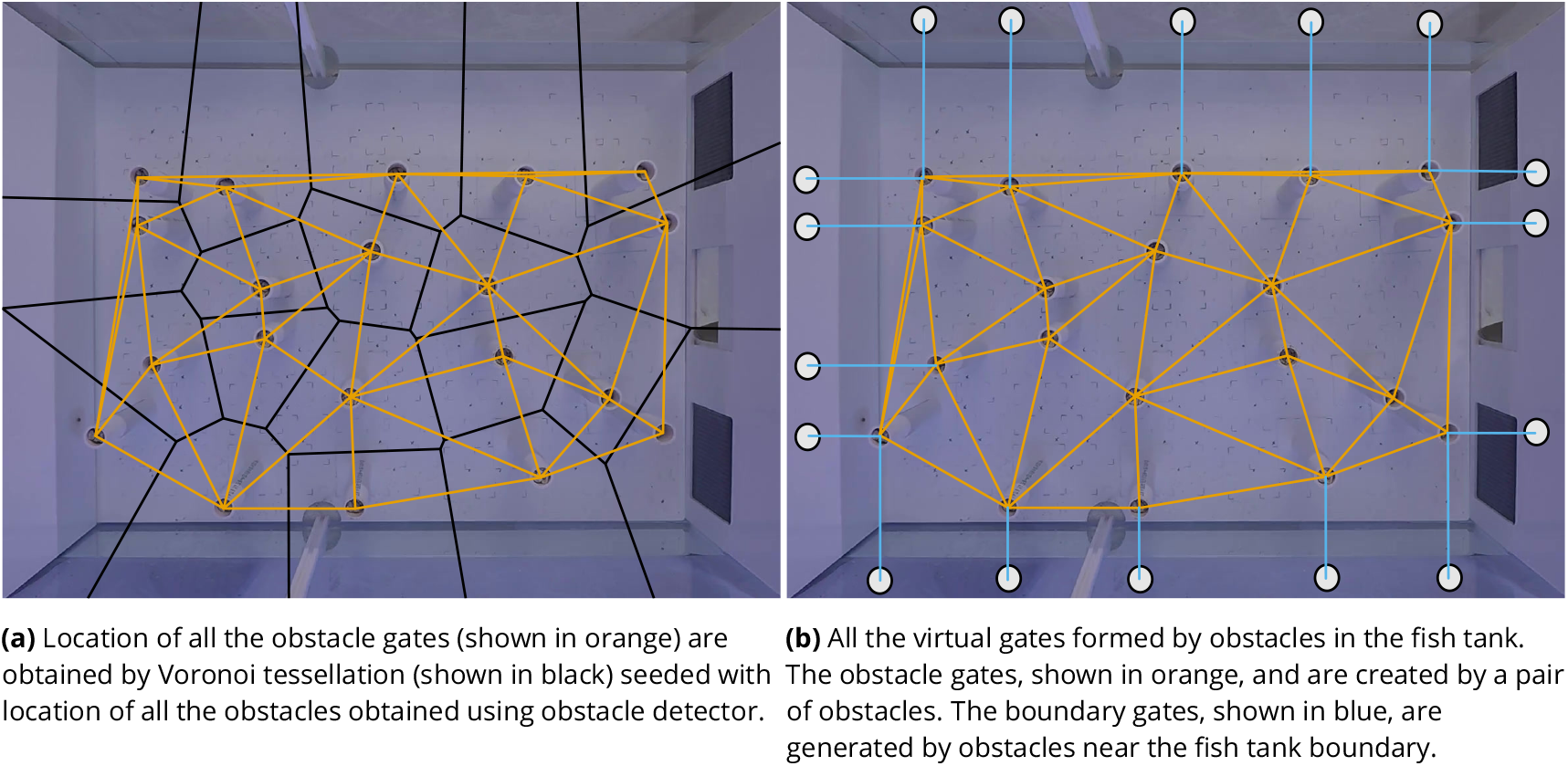
Computing the position of all the obstacles and virtual gates.

The process of locating the virtual gates between the obstacles and the aquarium walls is more challenging. These boundary gates are formed between the fish tank boundary and the obstacles near the boundary. The task of finding the location of boundary gates can be simplified if an imaginary obstacle is placed at the tank boundary, by translating the obstacles near the boundary while keeping them perpendicular to the tank surface as shown in Figure 9 (see top part). To accurately translate the obstacles to the boundary, two questions need to be answered:

**Figure 9.**
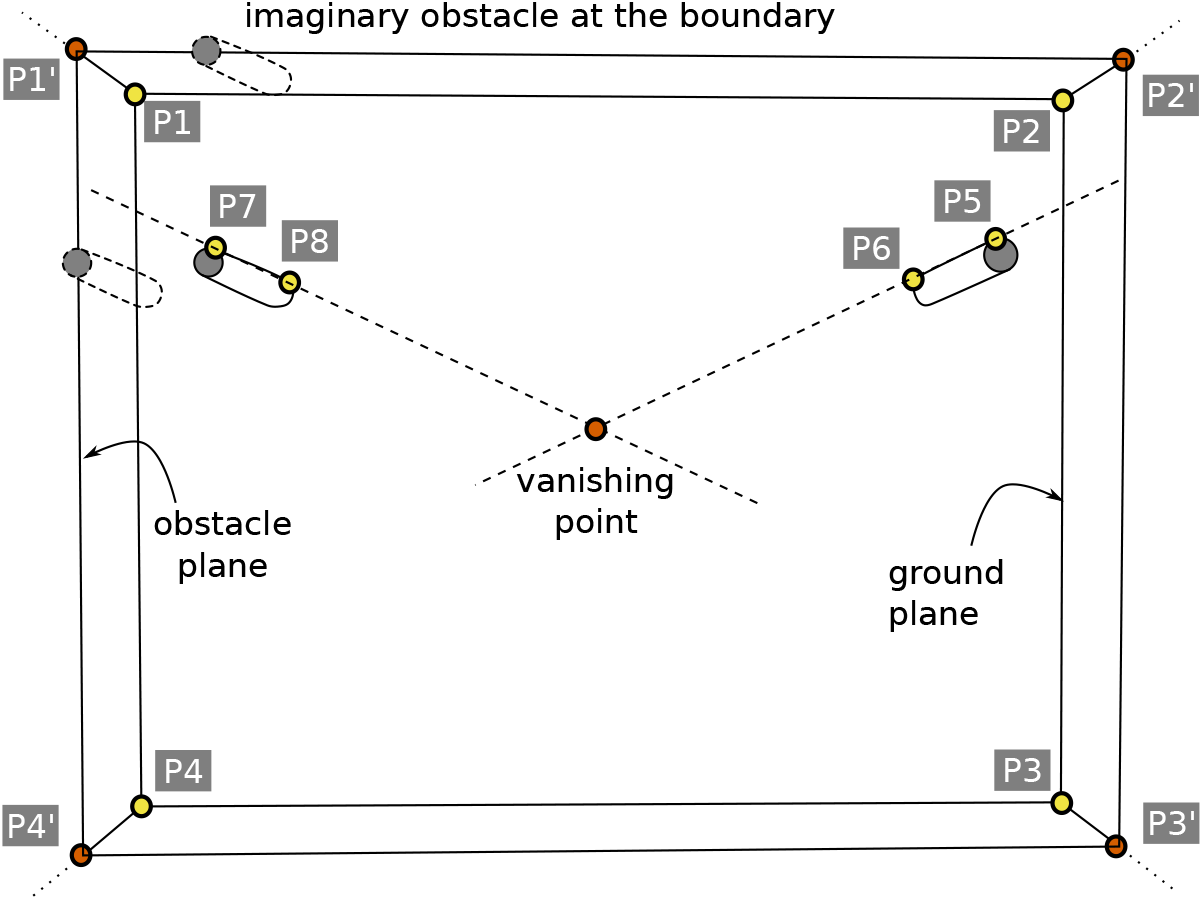
Four corners (P1, P2, P3 and P4) of the fish tank submerged under water are projected to the plane containing top part of all the obstacles (*i*.*e*. obstacle plane) using the homology formulation of (1). The projected corner points (P1′, P2′, P3′ and P4′) allows translation of obstacles to the fish tank boundary which helps form the boundary gates.

#### Box 1.

**Virtual gates Formed by the Obstacles**

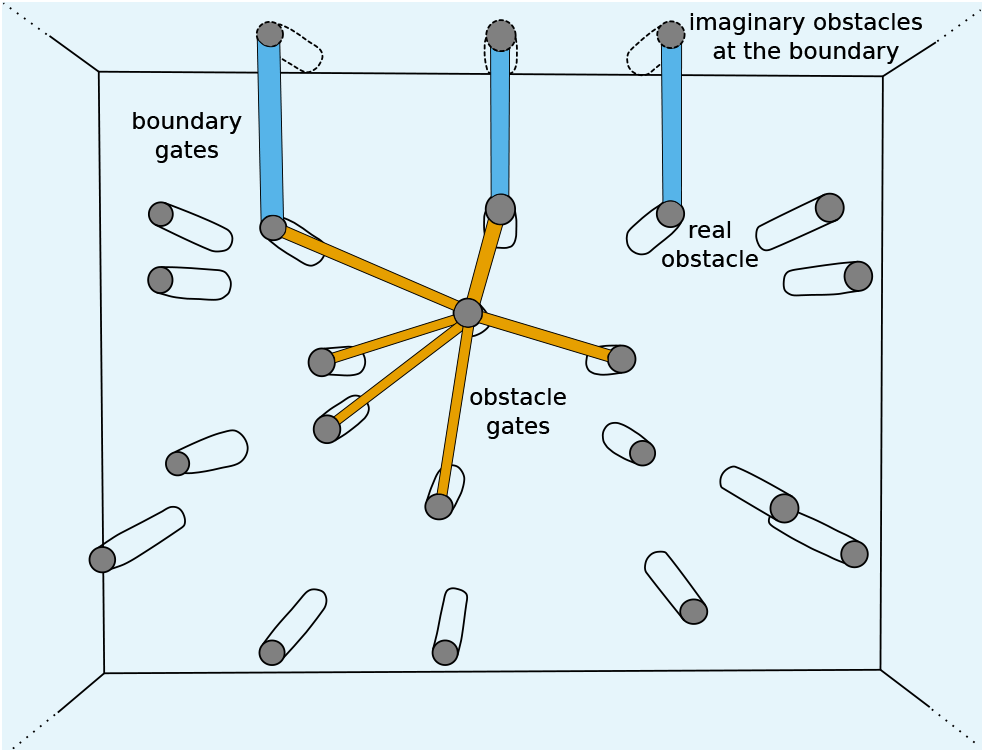
Virtual gates are formed by a pair of obstacles or by an obstacle near the tank boundary. The virtual gates can also be viewed as crossings through which the fish may decide to pass before reaching the food. An obstacle gate (shown in orange) is formed by a pair of obstacles while a boundary gate (shown in blue) is formed by an obstacle near the fish tank boundary. A fish crosses these gates during its journey to the food target.

1. Which obstacles are near the fish tank boundary?
2. Where is the fish tank boundary?

The first question is answered based on the Voronoi tessellation shown in Figure 8a (black). The Voronoi cells near the boundary have one of their boundary points extending towards infinity (i.e. beyond the edges of the image). All the cells can be computationally inspected to find the ones whose boundary points lies beyond the image boundary thereby revealing all the obstacles that lie near the tank boundary.

The second question appears trivial but requires significant effort to answer. It is difficult to automatically detect the four corners (or the boundary) of the fish tank as there are no clearly distinguishable visual features that can be used by automatic detectors. Therefore, help from human annotators is required. The calibration process described in the Calibration section (below) provides all the required feature points. The term ground plane denotes the plane containing the four corners of the bottom of the fish tank. All the cylindrical obstacles have the same length and therefore a plane passes through the top part of all the obstacles. The term obstacle plane denotes the plane containing the top part of all the obstacles. The tank corners are defined for the ground plane, whereas the coordinates of the obstacles (top part) are contained in the obstacle plane. Therefore, it is not possible to simply translate the obstacles to the tank boundary using the four corners manually defined in the ground plane. In order to translate the obstacles to the tank boundary using the coordinates of the obstacles (top part), the four manually defined corners (P1, P2, P3, P4) of the fish tank need to be projected from the ground plane to the obstacle plane.

The homology of parallel planes, described in ***Criminisi et al. (2000***), is used to project points from one plane to another. If *x* and *x*′ are a pair of corresponding points in the ground plane and obstacle plane respectively, then the two points are related as:

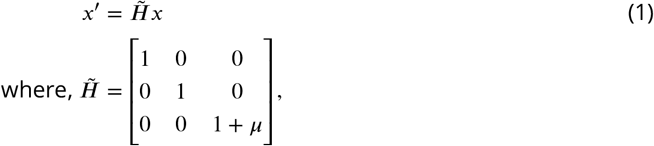

defines the homology for the case when the origin of image lies at the vanishing point with 𝜇 being the scale factor. The vanishing point is obtained by extending lines joining the feature points P5,P6 and P7,P8 as shown in Figure 9. These feature points were also obtained during the manual calibration process (see the Calibration Section). One pair of correspondences between the ground plane and the obstacle plane (*e*.*g*. P5 and P6) can be used to estimate the scale factor (𝜇). Using the homology 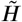, all the four tank corners (P1, P2, P3 and P4) are projected from ground plane to the obstacle plane (P1′, P2′, P3′ and P4′). The ground plane corner points are shown in yellow and the corresponding projected points are shown in orange in Figure 9.

The obstacles lying near the boundary are translated to the boundary as follows. For each boundary obstacle, the boundary edges (*i*.*e*. P1′P2′, P2′P3′, …) that intersect with the Voronoi cell boundary containing that obstacle are found. A Voronoi cell may intersect with more than one edge. For example, the obstacle containing manually annotated points P7 and P8 is contained in a Voronoi cell that intersects with the following two tank boundary edges: P1′P2′ and P1′P4′. After determining the boundary edges that intersect the Voronoi cell, the obstacle containing that Voronoi cell is translated to all the intersecting edges. This involves simply translating the real obstacle (top part) coordinates to the fish tank boundary (in the obstacle plane) as shown in Figure 9. Without the projected boundary (i.e. P1′, P2′, P3′ and P4′), it would not have been possible to perform this translation of the obstacles.

All the virtual gates computed using the process described above are shown in Figure 8b.

### Motion Analysis

The behavioural experiment described in this paper aims to understand the motion planning capabilities of Picasso triggerfish, which requires answering the following questions for each experimental trial video:

1. What is the length of virtual gates formed by the obstacles (*i*.*e*. the size of the gap between the obstacles), and which of the these gates are crossed by the fish during its journey to the food?
2. What is the speed, motion direction and body centroid of the fish during its journey to the food?

The first question is answered using the fish trajectory points computed in Section Detect+Track: Tracking High-Confidence Fish Detections and the location of virtual gates computed in Section Locating virtual gates Formed by the Obstacles. The fish is said to cross a virtual gate if one of the points in the fish trajectory lies within the bounds of the virtual gate. The third sub-figure in Figure 2 shows the fish trajectory (as a black line) and all the virtual gates crossed by the trajectory as green regions.

The second question is answered by estimating the apparent motion of pixels, within the fish bounding box, in consecutive frames. A close observation of two consecutive video frames, as shown in Figure 10 (top row), reveals that only the region occupied by the fish body changes between the two consecutive frames. Since the motion between two consecutive frames is small and corresponds solely to the fish body, the dense optical flow formulation implemented in the OpenCV software library is used to estimate the flow field of the fish motion as shown in Figure 10 (bottom left). The flow field also delineates (or segments) the pixels occupied by the fish body. Therefore, we use the flow field – instead of the rectangular bounding box around the fish body – to estimate the fish body centroid because the flow field only includes the regions occupied by the fish body and not the background regions near the fish body. This results is more accurate estimation of fish body centroid particularly when the fish body is curved (e.g. a c-shape).

**Figure 10.**
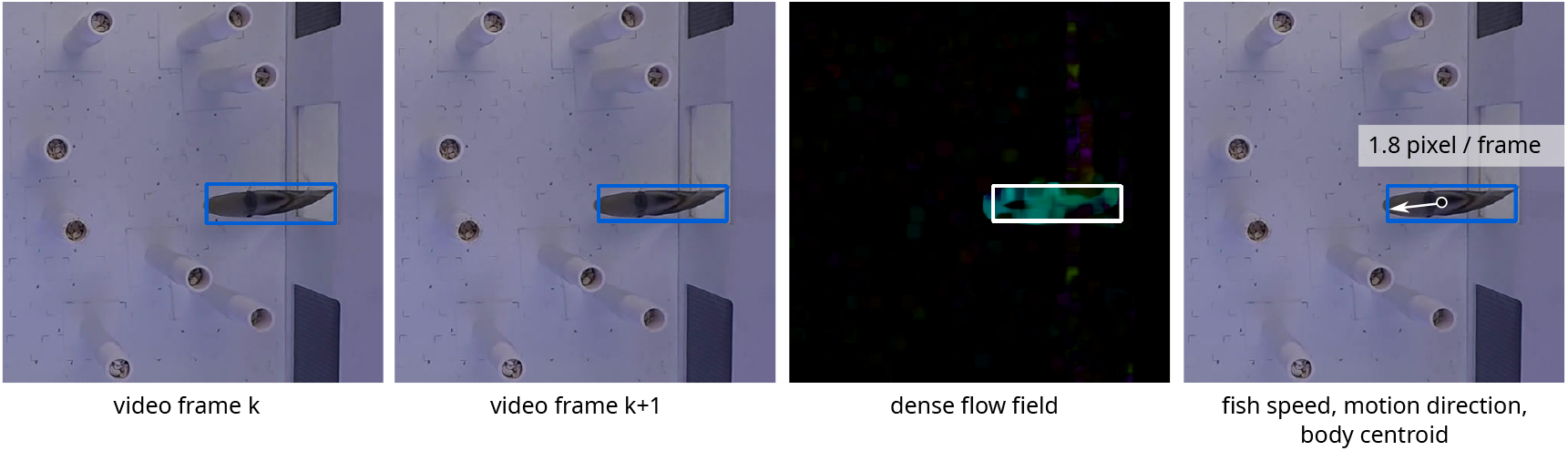
Motion of fish between consecutive video frames (top row) is estimated using optical flow. The dense flow field shows the optical flow magnitude. The estimated motion of each pixel reveals fish speed, motion direction and body centroid.

The optical flow formulation results in a 2-dimensional vector (*e*.*g*. [2.5, 3.1]) for each pixel location as shown in Figure 11 (top). These vectors capture the direction and speed of pixels in consecutive frames. Motion analysis is restricted to the pixel locations lying within the bounding box [*x*_0_, *y*_0_, *x*_1_, *y*_1_] describing the fish location. To help with visualisation of the flow field, this figure uses long vector arrows to represent pixels moving with higher speed and shorter vector arrows to represent pixels with lower speed. The two dimensional motion vector at pixel location (*x, y*) is denoted by *f* (*x, y*) = [*f*_*x*_, *f*_*y*_]. The magnitude of this vector is computed as 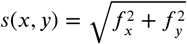 and this scalar denotes the speed of the pixel at location (*x, y*) as shown in Figure 11 (bottom). The pixels corresponding to the fish body have a higher speed while other pixels have nearly zero speed because only the regions corresponding to the fish body have apparent motion between consecutive frames. To mask out the regions that do not correspond to the fish body, we create a mask field *m*(*x, y*) whose value is 0 for all spatial locations where the pixel speed *s*(*x, y*) is less than a threshold (*e*.*g. t* = 0.1) and the value is 1 otherwise. Such a binary mask *m*(*x, y*) allows us to compute fish body centroid (*c*_*x*_, *c*_*y*_) as follows:

**Figure 11.**
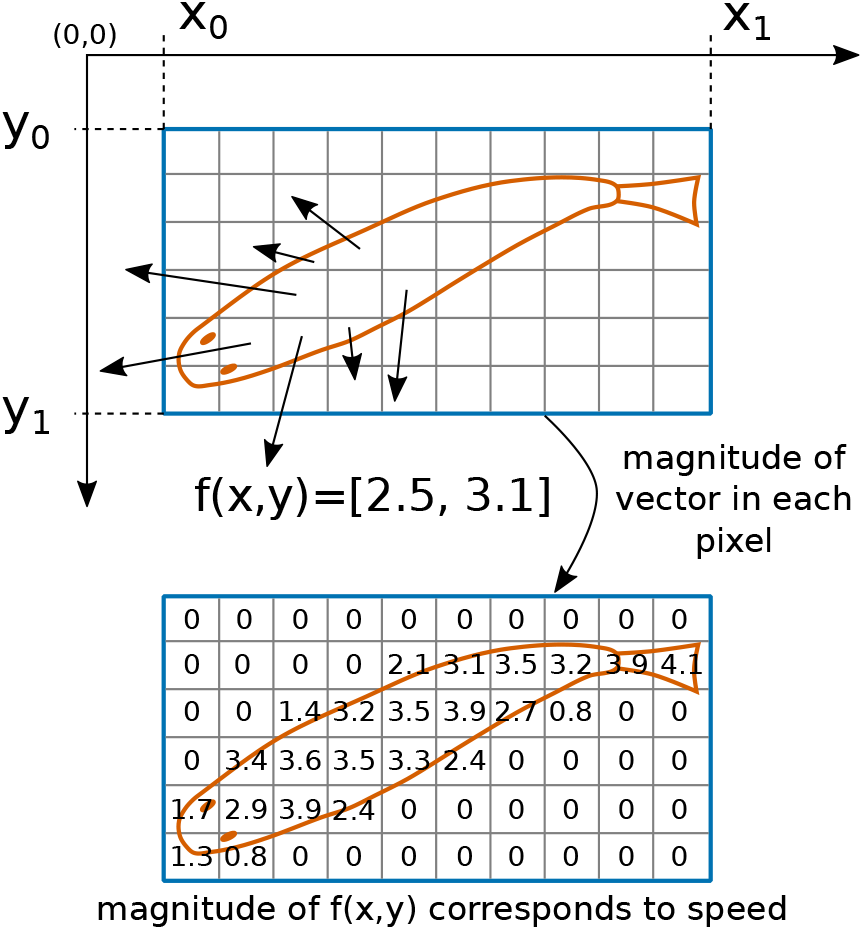
The optical flow field *f* (*x, y*) = [*f*_*x*_, *f*_*y*_] in the region containing the fish is used to compute the speed, motion direction and body centroid of the fish. Magnitude of the flow field vector (*i*.*e. s*(*x, y*)) denotes the apparent speed of piels to which a threshold is applied to mask out regions that do not correspond to the fish body.

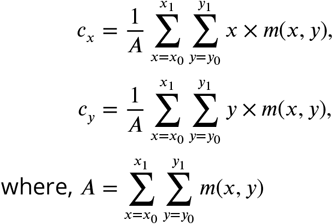

denotes the total area of the fish body mask. The computed fish body centroids are joined together to form the fish trajectory. Hence, the computation of the fish body centroid is dependent on the fish bounding box computed by the detector and tracker combination as well as on the optical flow field computed within the bounding box across two adjacent frames.

The motion direction of the fish is denoted by the 2-dimensional vector *v* which is computed by averaging only the flow vectors that lie within the fish body mask as follows

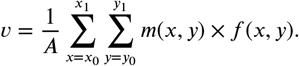

### Calibration

The vantage point of the overhead camera results in a view as shown in Figure 12. The automatic video processing method described in this paper requires manual annotation of 8 calibration feature points to locate the gates (or crossings) formed the fish tank boundary and the obstacles near the boundary. The first four calibration feature points (*i*.*e*. P1, P2, P3 and P4) correspond to the four corners of the fish tank boundary. The next four calibration points (*i*.*e*. P5, P6 and P7, P8) are placed on a pair of parallel vertical planes. These calibration points are used in Section Locating virtual gates Formed by the Obstacles to locate the intersection point of vertical parallel planes (*i*.*e*. the vanishing point) and create imaginary obstacles at the boundary that mirror the obstacles near the boundary. The imaginary obstacles at the boundary are required to automatically locate the gates near the boundary.

**Figure 12.**
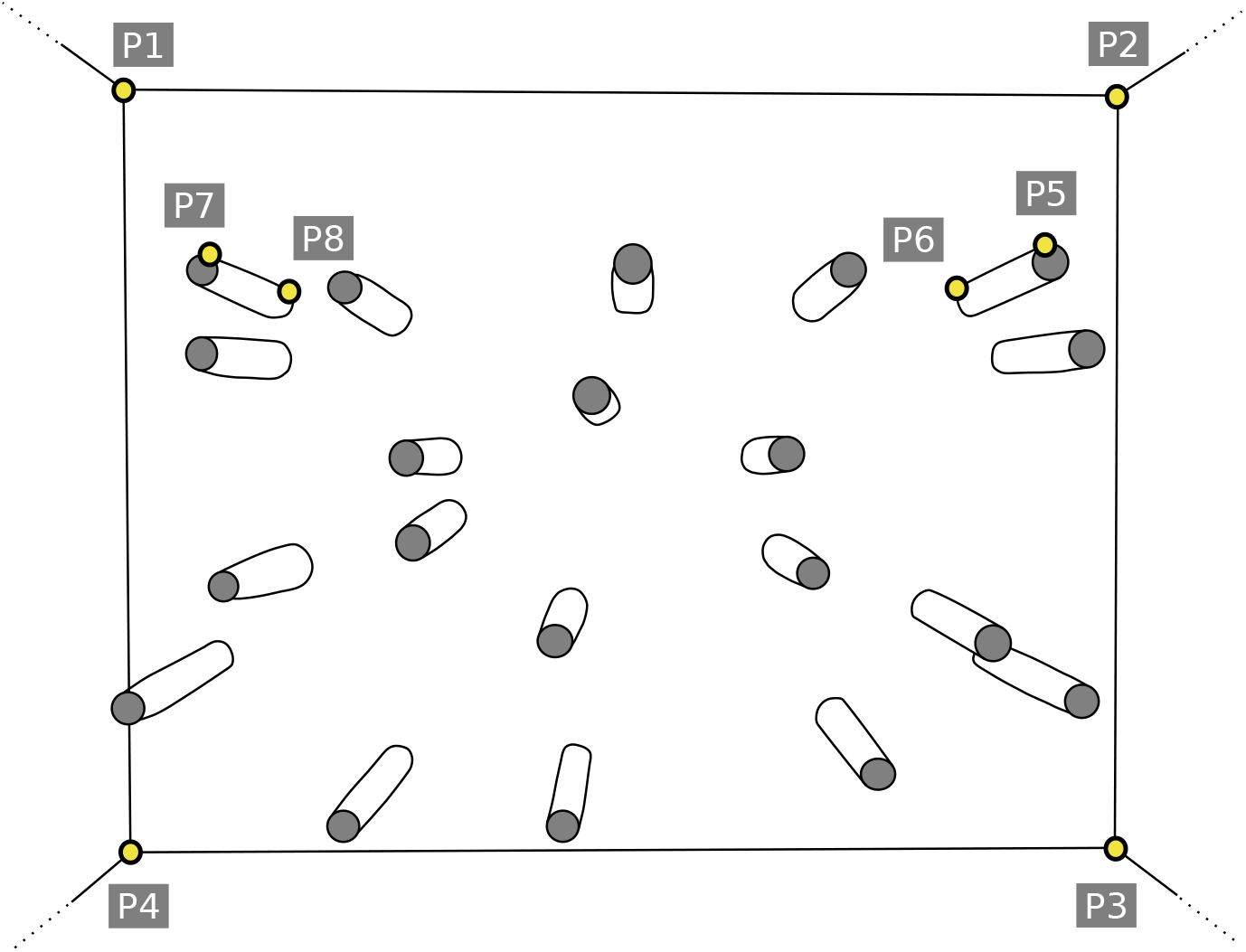
Initial calibration of the camera’s visual field involves manually annotating of the following 8 calibration feature points: four corners of fish tank boundary (P1,P2,P3 and P4), and feature points to locate top and bottom points on a pair of obstacles (P5,P6 and P7,P8). These calibration points are essential to accurately locate the virtual gates formed by obstacles.

The calibration process is one-off and has to be done only for any new setup of the camera or fish tank. We have provided the calibration data for all videos in our dataset. For any new setup, new calibration data can easily be obtained by manually annotating the 8 calibration feature points using our custom manual annotation tool included as a part of this work. In our experiment, the position of the video camera and the fish tank does not change during different experimental trials, therefore the calibration is only repeated where the camera, tank position, or camera field of view changes.

## Acknowledgments

CN was funded through a Leverhulme Trust Early Career Fellowship. AD and AZ are funded by the Visual AI: An Open World Interpretable Visual Transformer (UKRI Programme Grant EP/T028572/1). NP-C was supported by the Biotechnology and Biological Sciences Research Council (BBSRC) grant (BB/M011224/1).

## Authors’ Contributions

Authors CN, NP-C and GT designed the behavioural study. NP-C and CN carried out behavioural experiments. AD and AZ developed the software and tutorial, and CN tested it and provided comments. AD and CN drafted the manuscript and all authors commented on drafts.

## Data Accessibility

The tools and data used to evaluate the program are freely-available at https://www.robots.ox.ac.uk/~vgg/research/fish

## Footnotes

1 https://oecd.ai/en/catalogue/metrics/percentage-of-correct-keypoints-pck

2 Throughout this paper, the term high confidence detection refers to detections with confidence > 70%

3 https://gitlab.com/vgg/fish-tank-obstacles/-/tree/master/track/siamrpn

